# Non-rodent mammalian zygotes assemble dual spindles despite the presence of paternal centrosomes

**DOI:** 10.1101/2020.10.16.342154

**Authors:** Isabell Schneider, Marta de Ruijter-Villani, M. Julius Hossain, Tom A. E. Stout, Jan Ellenberg

**Affiliations:** Cell Biology and Biophysics Unit, European Molecular Biology Laboratory, Meyerhofstraße 1, 69117 Heidelberg, Germany; Department of Clinical Sciences, Faculty of Veterinary Medicine, Utrecht University, Yalelaan 112, 3584 CM Utrecht, the Netherlands

## Abstract

The first mitosis of the mammalian embryo must partition the parental genomes contained in two pronuclei. In rodent zygotes, sperm centrosomes are degraded and, instead, acentriolar microtubule organizing centers and microtubule self-organization guide the assembly of two separate spindles around the genomes. In non-rodent mammals, including human or bovine, centrosomes are inherited from the sperm and have been widely assumed to be active. Whether non-rodent zygotes assemble a single centrosomal spindle around both genomes, or follow the dual spindle self-assembly pathway is unclear. To address this, we investigated spindle assembly in bovine zygotes by systematic immunofluorescence and real-time light-sheet microscopy. We show that two independent spindles form around the parental genomes despite the presence of centrosomes, which had little effect on spindle structure and were only loosely connected to the two spindles. We conclude that the dual spindle assembly pathway is conserved in non-rodent mammals. This could explain whole parental genome loss frequently observed in blastomeres of human IVF embryos.

**Summary:** This study investigates spindle assembly during the first embryonic division in bovine zygotes that, like human, inherit centrosomes from the sperm. It shows that two independent microtubule arrays form by self-organization around parental genomes with only loosely connected centrosomes.

## Introduction

Mammalian fertilization involves the fusion of a sperm cell with an oocyte to give rise to a totipotent zygote, from which a whole new organism can develop. Faithful first mitotic divisions are essential for early embryonic development and to establish a healthy pregnancy. However, instead of being highly safeguarded, the first divisions in human embryos are surprisingly prone to chromosome mis-segregations and thus often lead to aneuploidy (Wells and Delhanty, 2000; Vanneste et al., 2009; Mertzanidou et al., 2013; Vera-Rodriguez et al., 2015; Munné et al., 2017; Fragouli et al., 2017). Postzygotic, or “mosaic”, aneuploidy, where a subset of cells in the embryo has an aberrant number of chromosomes, has been reported in up to two thirds of early human embryos produced *in vitro* (Wells and Delhanty, 2000; Daphnis et al., 2008; Vanneste et al., 2009; van Echten-Arends et al., 2011; Taylor et al., 2014; Vera-Rodriguez and Rubio, 2017). The high incidence of aneuploidy within the embryo is a major cause of developmental failure and pregnancy loss; and embryonic mosaicism is a major obstacle for embryo assessment after *in vitro* fertilization (IVF) in fertility clinics (Vanneste et al., 2009; Taylor et al., 2014; Munné et al., 2017; Fragouli et al., 2017; Vera-Rodriguez and Rubio, 2017). A similarly high degree of postzygotic aneuploidy has been reported in porcine, nonhuman primate, murine, bovine and equine embryos, suggesting that this phenomenon is common in the preimplantation development of many mammalian species (Zudova et al., 2003; Dupont et al., 2010; Bolton et al., 2016; Tšuiko et al., 2017; Shilton et al., 2020). Despite the widespread occurrence and often severe developmental consequences of post-zygotic aneuploidy, due to limited access to the relevant samples and technological difficulties to visualize these events in live mammalian embryos, we do not yet understand why the critical cell divisions at the beginning of mammalian life are so error prone.

The first division of the embryo is an exceptional mitosis. After fertilization, the parental genomes are replicated within the two separate pronuclei (PNs). Upon entry into mitosis, the nuclear envelopes break down and the two spatially separated sets of parental chromosomes have to interact in a coordinated fashion with the assembling mitotic apparatus of the zygote to allow synchronous and faithful segregation into two daughter cells. It was long assumed that the parental genomes would mix immediately after nuclear envelope breakdown (NEBD) and subsequently be segregated using a single zygotic spindle. In fact, even the definition of when a fertilized oocyte becomes a human embryo is based on the time when the parental genomes merge in some legal systems (e.g. Germany, § 8 Abs. 1, Embryonenschutzgesetz). However, using high resolution imaging of live embryos by light-sheet microscopy, we recently showed that, in mouse zygotes, two separate microtubule arrays form around each of the two parental genomes and keep the two genomes separated throughout the first mitotic division (Reichmann et al., 2018b). These two bipolar spindles first assemble and congress the parental chromosome sets independently in pro-metaphase. Then in metaphase, they align their pole-to-pole axes in order to segregate the two chromosome sets in parallel during anaphase. However, if the alignment of the two spindles is perturbed, the parental genomes can be segregated in different directions, leading to gross mitotic aberrations (e.g. formation of binucleated blastomeres or direct cleavage to 3 or 4 daughter cells) reminiscent of clinical phenotypes observed in human IVF embryos (Reichmann et al., 2018b).

Unlike most mammalian species, rodent zygotes do not contain centrosomes, with the sperm centrioles appearing to degenerate completely during spermiogenesis (Manandhar et al., 1998; Woolley and Fawcett, 1973). Instead, numerous acentriolar cytoplasmic microtubule organizing centers (MTOCs) are present during the first divisions, and the assembly of the bipolar spindles relies on microtubule self-organization and MTOC clustering (Courtois et al., 2012; Reichmann et al., 2018b). By contrast, nonrodent mammalian zygotes, such as human, porcine or bovine, inherit the centrioles from the sperm. Thus, in principle, they have two centers of cytoplasmic microtubule nucleation from the onset of mitosis (Fishman et al., 2018; Sathananthan et al., 1996). However, it is not clear whether these centrioles are in fact fully functional and how spindle assembly in these species proceeds. It might proceed analogously to that in somatic cells, where two centrosomes are the dominant centers of microtubule nucleation and also ensure formation of a single bipolar array early in mitosis. Alternatively, the mechanism may be similar to the mouse zygote involving the self-assembly of two separate bipolar arrays. While on the one hand human IVF phenotypes would suggest that the mechanism in nonrodents might be similar to that seen in the mouse, on the other hand the sperm centrioles have generally been assumed to be active (Fishman et al., 2018), which would argue for a single zygotic spindle. For obvious ethical and legal reasons, it was not possible for us to carry out high-resolution real-time imaging of spindle assembly using fluorescent markers in human embryos. We therefore decided to use the cow as a mammalian model organism to study how zygotic spindle assembly proceeds in the presence of paternal centrioles. As in human, bovine zygotes inherit the centrioles paternally, and both *in vivo* and *in vitro* produced pre-implantation cattle embryos show a high incidence of post-zygotic aneuploidies (Tšuiko et al., 2017). To study bovine zygotic spindle assembly, we combined systematic immunofluorescence (IF) of bovine zygotes, fixed at different stages of the cell cycle, with real time imaging of live zygotes by light-sheet microscopy during the first mitotic division. Our data clearly indicates that dual spindle assembly is a conserved mechanism, even when paternally inherited centrosomes are present.

## Results and Discussion

### Two separate zygotic spindles assemble in the presence of centrosomes

To investigate whether two spindles can form in a mammalian zygote, which inherited two centrioles paternally at fertilization, we analyzed spindle assembly following *in vitro* fertilization of bovine oocytes. First, we performed 3D IF microscopy of zygotes fixed at different stages of the first embryonic mitosis, and stained for pericentrosomal material, microtubules and DNA. In the majority of zygotes, the parental PNs were positioned adjacent to each other in prophase; in pro-metaphase we observed that two microtubule arrays had formed around them in close proximity (proximate spindles; Fig. 1A). Their longitudinal axes were mostly aligned during the later pro-metaphase stage and appeared to be fused during early metaphase. In the later mitotic stages, it was therefore often not possible to clearly distinguish between fused dual spindles and a single spindle. However, in an unexpectedly large number of zygotes, the PNs were further apart and the two spindles assembled at a large distance of ~30-65 μm (distant spindles; Fig. 1B). Such distant dual spindles were evident at all mitotic stages (Fig. 1B-C), including anaphase, and were thus functional for segregating chromosomes. Often, the timing of mitotic progression was asynchronous between the two parental PNs. This was especially evident at nuclear envelope breakdown, and, albeit more rarely, was also observed in later mitotic stages (Fig. S1B). The asynchrony suggests that the two PNs can not only set up two distinct spindles but can also independently regulate their cell cycle progression, even though they share a common cytoplasm. Across all mitotic stages we could clearly score dual spindles in almost 1/3 of the 178 fixed zygotes and in 33 we found distant dual spindles (Fig. 1C). This finding is in agreement with a recent paper from Brooks and colleagues, who observed that in 19 of the 49 bovine zygotes (38%) undergoing the first mitotic division, the two parental genomes failed to merge and thus segregated independently (Brooks et al., 2020). During metaphase, we observed that 17% of the zygotes showed two distant, but clearly bipolar microtubule systems (Fig. 1C, n = 12/72). Surprisingly, in most of these distant spindle pairs, pericentrosomal staining indicated that one pole of each spindle was associated with a centrosome (monocentrosomal spindles, Fig. S1C-D, n = 17/22), whereas only few distant dual spindles were acentrosomal (Fig. S1C–D, n = 3/22) or could not be scored due to poor pericentrosomal staining (Fig. S1D, n = 2/22). By comparison, proximate, closely aligned (or fused) spindles in metaphase mostly showed one centrosome at each of the two spindle poles (bicentrosomal contralateral spindles, Fig. S1C-D, n = 30/43), although we also observed monocentrosomal spindles (Fig. S1C-D, n = 7/43) and, in one case, a spindle with both centrosomes at the same pole (bicentrosomal ipsilateral spindle, Fig. S1C-D, n = 1/43).

**Figure 1:**
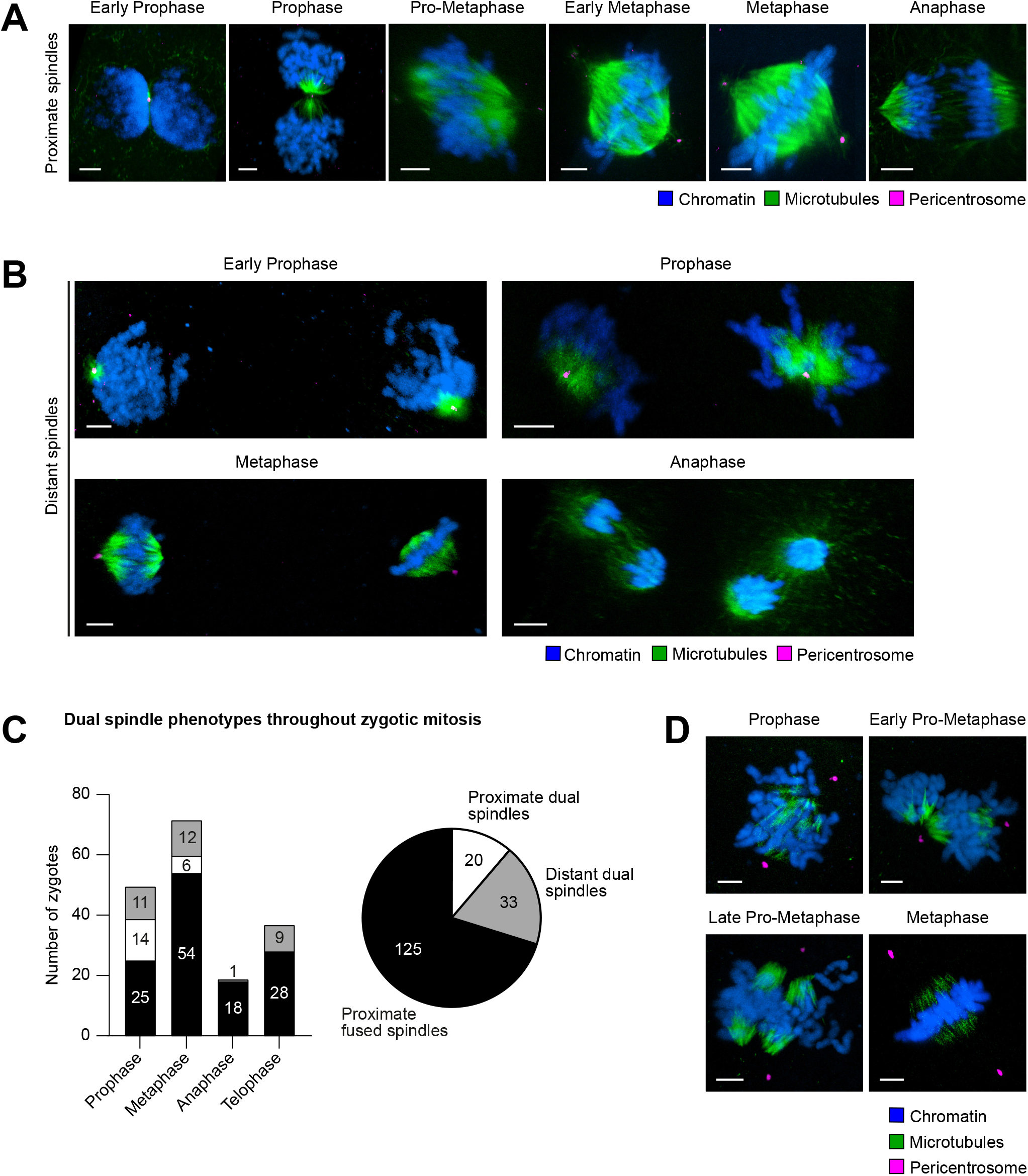
Dual spindle phenotypes in bovine zygotes. **(A and B)** Immunofluorescence of bovine zygotes fixed at 27.5 h post *in vitro* fertilization at consecutive stages of mitosis. Maximum intensity projections orthogonal to the estimated spindle axis of confocal sections showing proximate **(A)** or distant **(B)** dual spindles. Microtubules (alpha-tubulin, green); pericentrosomes (Cep192 or Nedd1, magenta); chromatin (Hoechst, blue). Scale bars, 5 μm. **(C)** Bar graph shows abundance of dual spindle types at different mitotic stages. Pie chart summarizes abundance of dual spindle types throughout mitosis. **(D)** Immunofluorescence staining of bovine zygotes (as in A), but following a cold shock on ice for 3 min prior to fixation. Maximum intensity projections of confocal sections orthogonal to the estimated spindle axis showing that centrosomal microtubules have been depolymerized beyond the detection limit. Microtubules (alpha-tubulin, green); pericentrosomes (Nedd1, magenta); chromatin (Hoechst, blue). Scale bars, 5 μm.

Together, these results demonstrate that dual spindle assembly, i.e. one around each of the two PNs, also occurs in mammalian zygotes that contain two centrosomes and, if the two PNs are distant, remains pronounced until chromosomes segregate. This also provides an explanation on how zygotes, inheriting centrosomes paternally, keep the parental genomes apart throughout the whole first mitotic division (Cavazza et al., 2020), similarly to mouse zygotes (Reichmann et al., 2018b).

Most commonly, both centrosomes localized to the opposite poles in proximate fused spindles, and in distant spindles each spindle showed one polar centrosome. Nonetheless, centrosome distribution varied, and we observed bipolar microtubule arrays that were able to segregate the chromosomes even if one or both poles lacked a centrosome. This suggests that the presence of centrosomes is not essential for spindle assembly and chromosome segregation in bovine zygotes. Whether distant monocentrosomal spindles are a consequence of incomplete pronuclear migration or abnormal pronuclei-centrosome interaction, remains to be determined.

### Centrosomes are only weakly linked to the spindle body

In both mono- and bicentrosomal spindles, the centrosomal microtubules appeared sparse and connected the centrosome to the body of the spindle only weakly. This was especially evident in fully assembled spindles from early metaphase onwards (Fig. 1A and B), and is in contrast to the canonical somatic spindle, where stable kinetochore fibers connect the centrosomes directly to the chromosomes (Prosser and Pelletier, 2017). To examine the strength of the connection between the centrosomal asters and the spindle body in the zygote, we subjected zygotes to a brief cold treatment to depolymerize unstable microtubules prior to fixation. Under these conditions, the microtubule bundles in the spindle body around the chromosomes were preserved, but the microtubules emanating from the centrosomes decreased to below the detection limit at all mitotic stages (Fig. 1D). After removing unstable microtubules in this manner, the gap between the spindle body and the centrosome increased significantly from 3.9 to 6.5 μm on average in metaphase (d_1_, Fig. S1E-F, p = 0.01), similar to the length of the half spindle after cold treatment (d_2_, Fig. S1E). Indeed, the ratio between the centrosome distance and the half-length increased from ~49% in unperturbed zygotes to 87% at cold treatment (d_1_/d_2_, Fig. S1G, p = 0.006). In addition, the centrosomes appeared to have moved somewhat away from the chromosomes, as their distance from the metaphase plate increased consistently, yet not significantly, after cold treatment (d_3_, Fig. S1E, H, p = 0.15). Additionally, we noted that in cold-treated zygotes, the two separate spindles forming around the parental genomes became more clearly visible, because a large gap had opened up between the remaining stable microtubule arrays as a result of the cold treatment (Fig. 1D, late pro-metaphase). This data demonstrates that the sparse microtubules connecting the centrosome to the spindle body as well as the microtubules between the dual spindles are unstable. This suggests that the centrosomes are only weakly linked to the spindle body and that the connection between the two spindles is also driven by dynamic microtubules.

### Centrosomes do not make a major contribution to metaphase spindle architecture

We next asked whether the weakly connected polar centrosomes influenced zygotic spindle architecture significantly. To answer this, we took advantage of the frequent occurrence of a monocentrosomal configuration in zygotes showing distant dual spindles (Fig. S1C-D). Although these separate spindles were smaller than proximate spindles, because they contained only one parental genome, they naturally offered the possibility to investigate whether the presence of a centrosome at only one pole induces an asymmetry between the spindle halves. As a control, we analyzed the degree of asymmetry in proximate fused spindles that had a centrosome at both poles. To measure symmetry, we computationally segmented the tubulin signal and quantified its spatial intensity distribution along the axis of the spindle orthogonal to the metaphase plate (Fig. 2A-B, S2A-B, for detailed description, see Materials and Methods). To compare the microtubule mass on both sides of the spindle, the total tubulin intensity within each spindle half – which corresponds to the area under the intensity distribution curve (AUC) of each half – was calculated and plotted ratiometrically (Fig. 2B-C). For monocentrosomal spindles, the total tubulin intensity in the centrosomal half of the spindle was slightly higher than that in the acentrosomal half (monocentrosomal, Fig. 2C, mean ratio = 1.2). This slight asymmetry was very similar to that between bicentrosomal spindle halves, when comparing the brighter to the dimmer half (bicentrosomal, Fig. 2C, mean ratio = 1.2). This indicates that a polar centrosome does not increase the microtubule mass in the associated spindle half by more than 20%, which is indistinguishable from the normal variation in microtubule mass between the halves of a bicentrosomal spindle (p=0.99).

**Figure 2:**
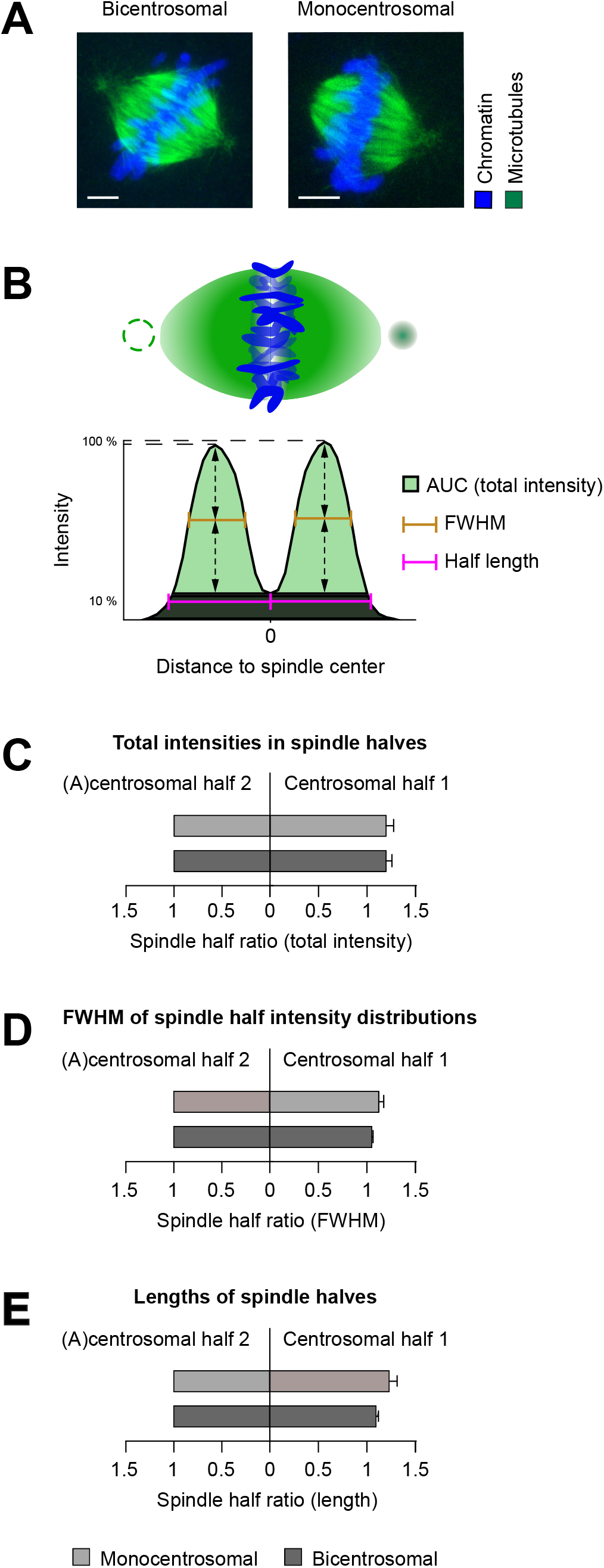
Quantitative comparison of proximate and distant dual spindles. **(A)** Exemplary immunofluorescence data subjected for quantitative comparison of proximate bi- and distant monocentrosomal spindles, see also Fig 1A-B. Metaphase spindles of bovine zygotes fixed at 27.5 h post *in vitro* fertilization. Maximum intensity projections over the imaging plane (z) are shown. (B) Schematic representation of zygotic metaphase spindle and total intensity distribution along determined spindle axis orthogonal to the chromosomes in bicentrosomal contralateral and monocentrosomal spindles. Note dashed circle to illustrate second centrosome in bicentrosomal spindles and missing centrosome in monocentrosomal spindles. Full width half maxium (FWHM) was considered as estimate of intensity distribution, area under the curve (AUC) as sum of total intensities in each spindle half and the half-lengths were calculated as distances between the intensity distribution’s valley (0 position) and the most distant positions along the axis, where the total intensity was 10% of the respective maximum. **(B-D)** Ratiometric comparison of intensity sum **(B)**, FWHM **(C)**, and of the length **(D)** between the halves of the spindle types. For distant monocentrosomal spindles (n = 11 from 6 embryos), absolute measurements were normalized to acentrosomal half. For proximate bicentrosomal contralateral spindles (n = 16), absolute measurements were normalized to the spindle half with lower sum intensity, or shorter FWHM and length. Bars indicate standard error of the mean. **(B)** Equal mean intensity ratio for the halves of mono- (1.21) and bicentrosomal (1.21) spindles (p = 0.99). **(C)** Comparable mean ratio of FWHM for the halves of mono- (1.14), and bicentrosomal (1.06) spindles (p = 0.12). **(D)** Overall similar mean ratio of spindle half lengths in mono- (1.24) and bicentrosomal (1.11) spindles (p = 0.07). Statistical test: unpaired t-test.

Even though the presence of a centrosome does not significantly change the amount of tubulin, it could still broaden its spatial distribution along the spindle axis away from the chromosomes/ equator. To investigate such subtle changes in spindle architecture, we measured the full width half maximum (FWHM) of the intensity distribution on each side of the metaphase plate for mono- and bicentrosomal spindles and again compared them ratiometrically (Fig. 2B, D). In monocentrosomal spindle halves, the FWHM increased slightly on the centrosomal side (monocentrosomal, Fig. 2D, mean ratio = 1.1), but the difference was not significantly different from that in bicentrosomal spindles when comparing brighter with the dim halves (bicentrosomal, Fig. 2D, mean ratio = 1.1; p=0.12).This indicates that a polar centrosome does not broaden the extension of dense tubulin away from the metaphase plate by more than 10%, indistinguishable from the normal variation found in bicentrosomal spindles. To investigate whether a centrosome might increase spindle length, for example by stabilizing microtubule bundles, we also compared the lengths of the halves within mono- and bicentrosomal spindles (Fig. 2B, E). In monocentrosomal spindles, we observed few asymmetric spindles, but on average the centrosomal half was only slightly longer than the acentrosomal half (monocentrosomal, Fig. 2E, mean ratio = 1.2). This difference was not significantly different from the length asymmetry observed in bicentrosomal spindles (bicentrosomal, Fig. 2E, mean ratio = 1.1, p=0.07). In summary, our quantitative analysis of the two halves of mono- or bicentrosomal spindles demonstrates that the presence of a centrosome at one pole of a zygotic metaphase spindle does not lead to a significant asymmetry in spindle structure, regarding either the total amount or the spatial distribution of microtubule mass, or the spindle half-length. Combined with the finding that centrosomes are only weakly linked to the spindle body and that acentrosomal spindles could segregate chromosomes, we conclude that centrosomes do not make a major contribution to the structure of the metaphase spindle in bovine zygotes nor are they required to initiate mitotic exit.

This is in contrast to somatic cells, as for example reported recently in centrinone-treated RPE1 cells, where centriolar halves are ~50-70% longer than acentrosomal halves (Dudka et al., 2019). The surprising symmetry of acentrosomal and centrosomal spindle halves in the zygote suggests that centrosome-independent pathways play a more important role for spindle assembly and maintenance. Nevertheless, centrosomes may have other important functions in the zygote such as pronuclear migration and the recently reported chromosome clustering at the pronuclear interface (Cavazza et al., 2020).

### Real time imaging reveals the dynamic process of dual spindle assembly

Although we analyzed a large number of zygotes (1421, of which 178 were undergoing mitosis), it was difficult to infer the precise order of the dynamic steps of dual spindle assembly in the presence of paternal centrosomes from snapshots of individually fixed embryos, primarily because of poor synchronicity, and variability in pronuclear position. We therefore decided to visualize spindle assembly in real time in live bovine zygotes. Based on technology we development for *in toto* imaging of pre-implantation mouse embryos (Strnad et al., 2016; Reichmann et al., 2018a), we adapted the inverted lightsheet imaging pipeline for the larger and more strongly scattering bovine embryos (for details see Material and Methods). Using mRNA microinjection at the pronuclear stage (Jaffe and Terasaki, 2004), we transiently expressed live fluorescent markers for chromosomes (Histone 2B, H2B) and the growing tips or lattice of microtubules (End-binding protein 3, EB3, or Microtubule-associated protein 4, MAP4). The inverted and low dose light-sheet microscope allowed us to maintain IVF culture conditions for bovine embryos and image them in 3D with a high temporal resolution of 2.5 minutes throughout the first division. These novel real time data sets of bovine zygotic mitosis clearly demonstrated that, indeed, two microtubule arrays assembled around the parental genomes in the presence of two centrosomes. Live imaging of a total of 21 dividing embryos revealed several different modes by which the two assembling spindles incorporated the two centrosomes, explaining the generation of the very different centrosome distributions that we had observed in fixed zygotes.

Consistent with the observations in fixed embryos, asynchronous NEBD of the two PNs was very common (n = 19/21), with a delay between the leading and lagging pronucleus (PN) ranging from 2.5 to 7.5 min. Independent of synchronicity, microtubules often accumulated within the original pronuclear volumes and two small microtubule asters formed around the centrosomes. In most of the zygotes the parental PNs had come into close proximity before NEBD (n = 20/21). The centrosomes were also mostly in contact with the pronuclear surfaces (Fig. 3A-B and S2C-D). In general, how the centrosomal asters were then associating with the two spindles forming around the chromosomes largely depended on their original orientation respective to the PNs; Most commonly (~60%) both centrosomes were wedged in between the two nuclear envelopes and thus associated with both parental genomes. From here, they were usually incorporated into one pole of each of the two developing spindles, in a revealing dynamic process: Both asters initially associated with the spindle that formed around the ‘leading’ PN, undergoing NEBD first (e.g. PN1, Fig. 3A and S2C, 7.5 min). Once the second PN also initiated NEBD (PN2, Fig. 3A and S2C, 15 min), microtubules transiently accumulated around its chromosomes and a second (half-) spindle formed between one of the centrosomes and the second genome. In cases where the spindle orientation relative to the light-sheet allowed high resolution imaging of this step, we observed that the microtubules accumulating around the second genome pulled one centrosome away from the first spindle incorporating it instead into the second, initially often monopolar array (Fig. 3A and S2C, 15 and 20 min post NEBD; Suppl. Movie 1). Subsequently, the second spindle also became bipolar and simultaneously, the two spindles aligned their axes in parallel (see Suppl. Movie 1). Finally, we could recognize the fused dual spindle with an overall round appearance and broad poles that we had often observed in fixed embryos (compare Fig. 3A and S2C, 57.5 min with Fig. 1A, metaphase). Interestingly, also in live embryos, some of these fused proximal metaphase spindles still had polar centrosomes, which were positioned slightly off-center and were only weakly connected to the spindle body (Fig. 3A and S2C, 20 and 57.5 min post NEBD; see Suppl. Movie 1). Rarely, no dominant initial bipolar array was developed but instead two monopolar and monocentrosomal spindles formed around the two PNs, eventually combining into a bipolar array.

**Figure 3:**
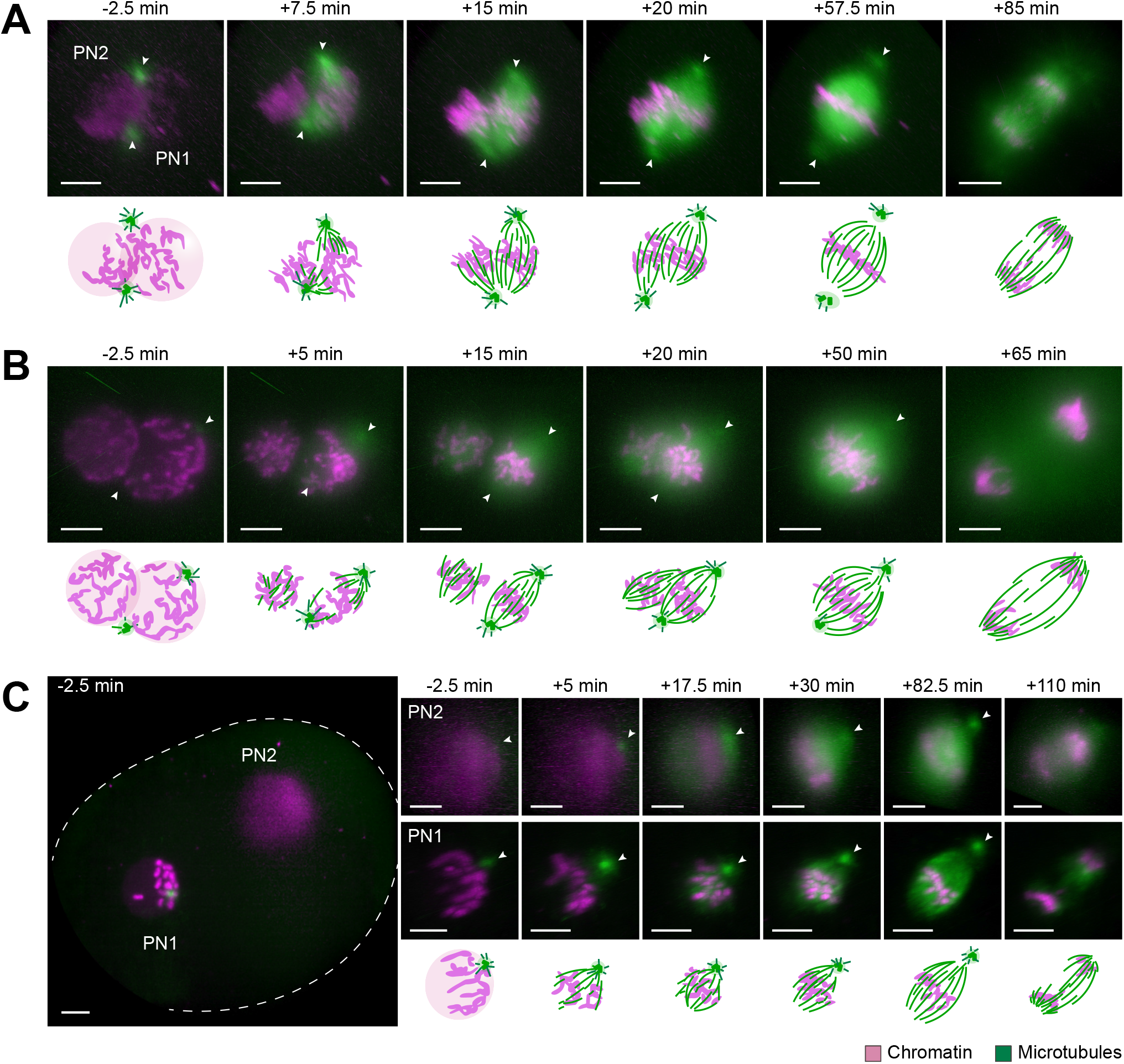
Assembly and dynamics of proximate and distant dual spindles in live bovine zygotes. **(A, B and C)** Bovine zygotes expressing microtubule markers (EGFP-MAP4, A) or EB3-mEGFP2, B and C; green) and chromatin marker (H2B-mCherry; magenta) were imaged by light sheet microscopy every 2.5 min throughout mitosis and for up to 6 h in total. 3D rendered images of pronuclear volumes are shown. Overview image to illustrate pronuclear distance (C) is a background corrected (median based denoising) overlay of maximum intensity projections over z of both pronuclear volumes within the zygote (zygotic rim indicated by dashed lines). Timings are respective to synchronous pronuclear envelope breakdown (NEBD) (B) or to NEBD of leading pronucleus (PN1) in case of asynchrony (A, C). PN2, lagging pronucleus. Arrow heads indicate positions of centrosomes. Projected scale bars, 10 μm. **(A)** Most frequent, and **(B)** most pronounced example of proximate dual spindle assembly. **(C)** Example of distant dual spindle assembly with two individual monocentrosomal spindles throughout mitosis.

In one striking example, we could distinguish the separate initial arrays over several minutes (Fig. 3B and S2D). Here, the two centrosomes were associated with opposite sides of only one PN and both centrosomes remained associated with the first spindle that formed around this PN. The second genome then clearly nucleated microtubules independently of centrosomes, forming a more spherical bipolar microtubule array. It first increased its microtubule mass before merging with the first, bicentrosomal array, pole by pole (Fig. 3B and S2D, 15-50 min post NEBD; see Suppl. Movie 2). Overall, in the embryos showing dual spindle assembly around adjacent PNs with closely associated centrosomes (Fig. 3A-B), the first mitosis usually resulted in a symmetrical two cell embryo (~85 %).

For the remaining zygotes with proximal PNs and associated centrosomes, dual spindle assembly was also evident (Fig. S3A-C). However, since the centrosomes were originally not located at opposite sites along the pronuclear interphase, the spindle configurations were more variable from embryo to embryo. Instead of assembling a dominant bipolar array, two spherical and/or monopolar spindles assembled around the two genomes seemingly independent of the centrosomes. In the initially forming spindles, centrosome positions ranged from polar but ipsilateral (Fig. S3A, >7.5 min post NEBD), to apolar (Fig. S3B, 10-22.5 min post NEBD). As mitosis progressed, the centrosomes were incorporated into different parts of the spindles. However, neither positioning ipsilaterally at one pole (Fig. S3A, 35 min post NEBD) nor at the spindle midzone (Fig. S3B, 55 min post NEBD) was corrected.

We also observed spindles with completely dissociated centrosomes (Fig. S3D-E, n = 3/21). Here, only one of the centrosomes was in the proximity of the PNs at NEBD, while the other was far away in the cytoplasm (Fig. S3D and E, 5 min post NEBD). Again, two microtubule arrays formed that eventually merged into a single bipolar spindle with no centrosome at one pole. If close enough, the second centrosome could be pulled in by the fully formed spindle (Fig. S3D, 25-45 min post NEBD), but if far away, it remained isolated in the cytoplasm (Fig. S3E). It would be interesting, in the future, to understand which intrinsic mechanisms could be responsible for the rather frequent phenomenon of centrosome displacement. It is possible that it results from an impaired connection of centrosomes to microtubules and the pronuclear envelope. Such disruptive connection could impair both, centrosome separation and movement.

One of the live imaged zygotes had its two PNs positioned approximately 60 μm apart. Strikingly, its two bipolar microtubule arrays remained completely separate until chromosome segregation (Fig. 3C and S2E, n = 1/21). Consistent with the fixed embryos (Fig. S1C-D), each PN of this live zygote was associated with one centrosome (Fig. 3C, 5 min, white arrowheads). After NEBD, initially two monocentrosomal spindles formed, which then bipolarized and progressed to chromosome segregation remaining over 50 μm apart (Fig. 3C and S2E, Suppl. Movies 3 and 4). This configuration did not result in a normal cleavage into a symmetrical two cell embryo, but exhibited several mitotic errors including ingression of multiple cleavage furrows and failure of cytokinesis (Fig. S2F, Suppl. Movie 5).

Together, our observations in living zygotes were fully consistent with the results obtained by IF and explained the temporal sequence of the spindle assembly intermediates we had observed in fixed zygotes. The real time data showed that two microtubule arrays with up to four ‘poles’ form around the two PNs, despite the presence of only two clearly astral microtubule organizing centers, i.e. the centrosomes. Moreover, they showed that in living zygotes centrosomes are not essential for bipolar spindle assembly and that their attachment to the spindle body is rather loose; for example, allowing the second spindle to capture and remove a centrosome from the first one, or a centrosome being associated to the spindle midzone rather than to the pole. They also showed that centrosomes were lost into the cytosol if localizing more than approximately 5 μm away from the PN or spindle body.

### Most spindle microtubules originate from the vicinity of chromosomes

In all live embryos (21/21), the centrosomes nucleated microtubules shortly before and at NEBD. Within 10 minutes after NEBD however, the bulk of spindle microtubules seemed to accumulate or even originate in the vicinity of the chromosomes; this was particularly evident when one of the two zygotic spindles was acentrosomal (Fig. 3B, left PN). Furthermore, centrosomal microtubules, when present, grew preferentially towards the DNA after NEBD and the microtubule signal intensities at centrosomes seemed to decrease already early in mitosis whereas microtubule mass at the spindle center seemed to increase quickly (Fig. 4A). To quantify where most of the spindle microtubule mass appeared at different times of early mitosis, we analyzed the changes in spatial distribution of total microtubule intensity over time along the centrosome axis, from prophase to pro-metaphase (Fig. 4B-D). We were interested in comparing the total microtubule mass (Fig. 4B, black cuboid with solid line), but also in comparing concentrations within equally small volumes along the centrosomal axis (Fig. 4B, black cuboid with dashed line). This analysis revealed that on average, within 5-7.5 min of NEBD, the total microtubule intensity started to increase at the spindle axis center (Fig. 4C, light dashed line; n=6), and reached a peak of ~85% by 20 min post NEBD. In comparison, we only observed a modest increase of less than 20% around the centrosomes (Fig. 4C; dark dashed lines), which peaked already around 5-7.5 min post NEBD. After this initial small increase, the centrosomal microtubule mass declined or stagnated, while the chromosomal microtubule mass continued to rise. Even when comparing the microtubule mass within equal volumes at the axis center (where chromosomes would be located; Fig. 4D, light dashed line) and at the centrosomes (Fig. 4D, dark dashed lines), we observed a similar behavior, indicating that microtubule concentration increases more in the vicinity of chromosomes. To analyze and visualize this change in microtubule abundance at the centrosomes and at the chromosomes further, we plotted the change in total mass (Fig. 4E) and in concentration (Fig. 4F) at both locations over time (n = 6). This analysis confirmed that microtubule mass only modestly and transiently increased at the centrosomes until 5-7.5 min post NEBD, whereas chromosomal microtubule mass continued to rise all the way to late prometaphase, when spindle assembly was largely complete. At metaphase, the spindles in live zygotes had a barrel shaped appearance, and mostly separated centrosomes (Fig. 3A-C; S2C-E, white arrow heads; Fig. 4A, pseudo color profile). Together, these results indicate that chromosomes most strongly contribute to microtubule nucleation and polymerization in the bovine zygote, and suggest that centrosomes contribute little to the increase in overall microtubule mass after NEBD. This is consistent with our observations of few and unstable centrosomal microtubules that make a weak connection to the spindle body in fixed and cold treated zygotes (Fig. 1D).

**Figure 4:**
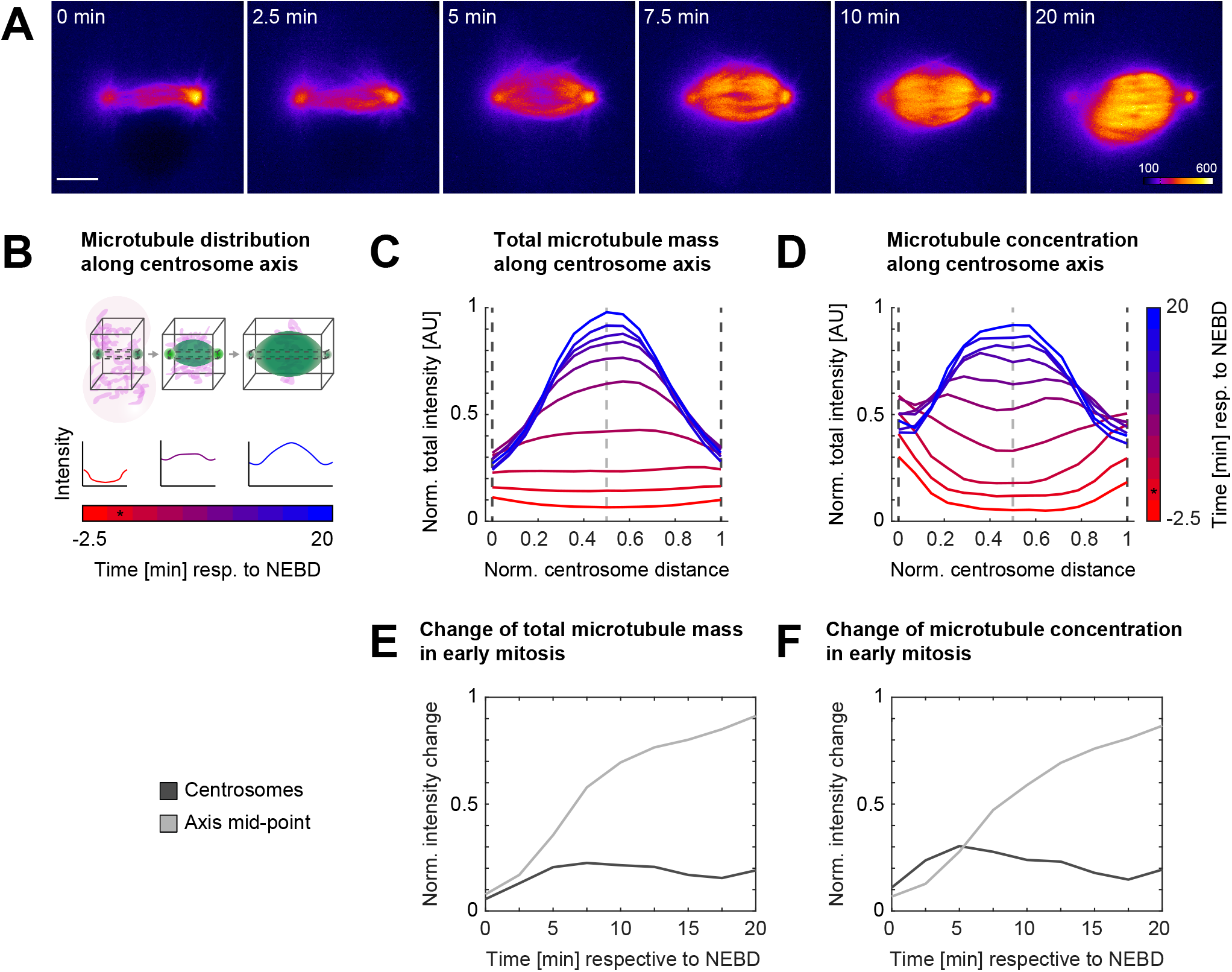
Analysis of dynamic distribution of spindle microtubules in live bovine zygotes. **(A-F)** Microtubule signal (EGFP-MAP4 or EB3-mEGFP2) from live imaging of bovine zygotes by light sheet microscopy every 2.5 min with a spindle assembly type as described in Fig. 3A was analyzed for 10 time points starting 2.5 min prior to nuclear envelope breakdown (NEBD) of the leading pronucleus (PN) or both PNs. **(A)** Pseudocolor representation of EGFP-MAP4 signal within single planes through the centers of intensities at centrosomes. Corresponding lookup table is depicted in last frame of the time series. 6 of 10 analyzed time frames were selected to visualize critical time points for microtubule redistribution in early spindle assembly. Time in min respective to NEBD. Scale bar, 10 μm. **(B-D)** Measuring intensity distribution of microtubule signal along the centrosomal axis over time.. **(B)** Scheme illustrating the measurements: After background subtraction, total microtubule intensities were calculated for 15 equidistantly distributed 2D slices within the black cuboids along the centrosomal axis for the different time points. Black cuboid with solid line encompasses 351×351 pixel sized slices to measure entire microtubule intensity or mass along the spindle axis. Small black cuboid with dashed line encompasses 15×15 pixel sized slices to indicate relative microtubule concentrations. Maximum normalized total intensities along the normalized centrosome distance were annotated. **(C-D)** Average distribution of maximum normalized microtubule intensities along centrosomal axis indicating **(C)** relative microtubule mass within spindles over time (black cuboid with solid line, as described in B), and **(D)** relative microtubule concentrations (black cuboid with dashed line, as described in B); n = 6 zygotes. Dashed lines mark the position of the 2D slice through the centrosomes (black dashed lines) and the centrosome axis mid-point (grey dashed line). Color gradient from red to blue indicates time in min respective to NEBD. Time of NEBD indicated by asterix. **(E-F)** Average change of normalized total microtubule intensity over time from NEBD until 20 min post NEBD, at centrosomes and the centrosome axis mid-point, indicated by black and gray dashed lines in C and D, respectively; n = 6 zygotes. For intensity change at centrosome, mean intensity of both centrosomes was calculated.

Overall, all our observations are consistent with a model where chromosomal microtubule nucleation and self-organization are dominant driving forces for bovine zygotic spindle assembly, while the weakly associated centrosomes make only a minor contribution. This explains why pronuclear position and timing of NEBD of the two PNs at the beginning of mitosis are the main determinants of how quickly the two spindles form, align and merge. It also explains why, when the two PNs are far apart, two spindles are generated that remain separate until chromosome segregation. The fact that spindles form by chromosomal microtubule self-organization, and incorporate the centrosomes only if they are within approximately 5 μm reach of their weak asters, also explains the very variable allocation of centrosomes to the four possible spindle poles, with all possible combinations within a bipolar microtubule array: from two centrosomes at one pole, to no centrosome at all. This is very different from the situation in somatic cells or *C. elegans* embryos (Prosser and Pelletier, 2017; Müller-Reichert et al., 2010), where the two centrosomes are the dominating centers of microtubule nucleation and thus, from the onset of mitosis, build a single bipolar microtubule array with well-focused poles that captures the chromosomes. Despite the presence of the two sperm centrioles, and eventually two centrosomes, the bovine zygote surprisingly behaves rather similarly to the mouse zygote where the sperm centrioles are degraded. The main difference being that, in the mouse, the two spindles cluster some of the many cytoplasmic MTOCs at their poles, whereas in the cow, the two centrosomes are incorporated seemingly randomly only if positioned close by. It will be very interesting in the future to understand the mechanism of chromosomal microtubule nucleation and spindle bipolarization in mouse and bovine zygotes and to carefully compare it to clinical data to infer whether a similar process occurs in human zygotes. Recent studies are pointing towards such a mechanism in human zygotes, especially the observation that the genomes frequently display a multipolar orientation in prometa- and metaphase and that chromosomes are frequently segregated in a uniparental conformation (Ford et al., 2020).

Different to the mouse, the larger bovine zygotes exhibited a striking degree of incomplete or failed pronuclear migration, which sometimes resulted in very large distances between the two PNs at the onset of mitosis. While this might be particularly prominent in *in vitro* fertilized zygotes, these distant PNs provide the opportunity to observe the formation of dual spindles over a longer time interval without alignment and merging. While the centrosomes do not seem to be essential for zygotic spindle assembly *per se*, they may play a role in coordinating pronuclear migration as has been reported in species such as *C. elegans* (Malone et al., 2003), and very recently also in bovine zygotes (Cavazza et al., 2020). In the future, it will be very interesting to investigate the pronuclear migration process in further detail and probe its robustness in mammalian zygotes. The similarities with human zygotes, such as the inheritance of the centrosomes from the sperm and the increased risk of mis-segregation during the early embryotic cleavages, make the bovine zygote a valuable model to study the mechanisms behind the error prone nature of early embryonic division in non-rodent mammals, and has important implications for improving the quality of infertility treatments and better understanding how the parental genomes in the embryo are partitioned and eventually merged.

## Materials and Methods

### Bovine oocyte collection, in vitro maturation, fertilization and zygote culture

Cumulus-oocyte complexes (COCs) were collected from abattoir ovaries as described by Ferraz and colleagues (Ferraz et al., 2018). Only COCs with a minimum of three layers of cumulus cells were selected and washed in HEPES-buffered M199 (22340-020, Gibco BRL, Paisley, U.K.) and then either directly matured in vitro for 23 hours in groups of 35–70 COCs in 500 μl of maturation medium (31100-027, NaHCO3-buffered M199 [11150059, Gibco BRL] supplemented with 1% (v/v) penicillin-streptomycin [15140122, Gibco BRL], 0.02 IU/ml FSH [Sioux Biochemical Inc., Sioux Centre, IA, USA], 0.02 LH IU/ml [Sioux Biochemical Inc.], 7.7 μg/ml cysteamine [30070, Sigma-Aldrich, St. Louis, Missouri, USA] and 10 ng/ml epidermal growth factor [E4127, Sigma-Aldrich]) at 38 °C in a humidified atmosphere at 5% CO_2_, or hold at room temperature for 19 hours in synthetic oviduct fluid for holding (H-SOF, Avantea, Italy) before in vitro maturation. After maturation the oocytes were fertilized using frozen thawed sperm cells from one bull of known fertility. Spermatozoa were selected by centrifugation through a discontinuous Percoll (90/45%; P1644, Sigma-Aldrich) gradient and added at a final concentration of 1 × 106 cells/ml to fertilization medium (Parrish et al., 1988) supplemented with 1.8 IU/ml heparin (H3393, Sigma-Aldrich), 20 μM d-penicillamine (P4875, Sigma-Aldrich), 10 μM hypotaurine (H1384, Sigma-Aldrich), and 1 μM epinephrine (E4250, Sigma-Aldrich). In vitro fertilization was performed for 6-9 h at 38 °C in a humidified atmosphere at 5% CO_2_. Presumptive zygotes were then vortexed for 3 min to remove cumulus cells and, transferred to synthetic oviductal fluid (SOF) (Takahashi and First, 1992) and cultured at 38 °C in a humidified atmosphere, at 5% CO_2_ and 5% O_2_. The zygotes used for live imaging were cultured in the same conditions with the absence of Phenol Red in the SOF culture media. On day 5 of culture, cleaved embryos were transferred to a new fresh SOF (500 μl per group of 35-70), and cultured further until day 8 under above described conditions.

### Immunofluorescence (IF) and confocal imaging

At 27.5 hours post fertilization, bovine zygotes were briefly washed in PBS at 38 °C and either directly fixed in 500μl fixation medium (94 mM PIPES pH 7.0, 0.94 mM MgCl_2_, 94 μM CaCl_2_, 0.1 % Triton X-100, 1 % PFA) or after incubation in ice cold PBS for 3 minutes (cold shock treatment) as described for mouse oocytes and embryos (Kitajima et al., 2011; Reichmann et al., 2018b). The embryos were then washed 4 times in 3% BSA in PBS with 0.1% Triton (PBS-T) at room temperature and extracted in PBS-T overnight at 4 °C. All the following treatments were done within wells of ibidi μ-Slides (81501, μ-Slide Angiogenesis, ibidi) filled with 40 μl of solution per well. Embryos were blocked in 5% normal goat serum and 3% BSA in PBS-T and then incubated with the primary antibodies in blocking solution overnight at 4 °C. The primary antibodies used were chicken anti alpha-tubulin (10 μg/ml; abcam, ab89984), rabbit anti CEP192 (3.5 μg/ml; Ab frontier AR07-PA001), mouse anti-acetylated tubulin (5 μg/ml; T7451 Sigma-Aldrich) or mouse anti NEDD1 (2.5 μg/ml; H00121441-M05, clone 7D10, Abnova). Embryos were then washed 3 x 5 min with 3% BSA in PBS-T and incubated with the following DNA dye and secondary antibody dilutions in blocking solution for 3 hours at room temperature: Hoechst 33342 (0.2 mM, Sigma-Aldrich), Goat Anti-Chicken Alexa Fluor® 647 (4 μg/ml, A-21449, Molecular Probes), Goat Anti-Mouse Alexa Fluor® 488 antibody (8 μg/ml, A-11029 Invitrogen), Goat anti Rabbit Alexa Fluor® 568 antibody (8 μg/ml, A-11036 Invitrogen). The embryos were then washed 3 x 10 minutes with 3% BSA in PBS-T and 2 x 10 minutes with PBS alone and mounted on glass slides (Superfrost Plus, Menzel, Braunschweig, Germany) with anti-fade mounting medium (Vectashield, Vector Laboratories, Burlingame, California, USA).

Fixed bovine zygotes were imaged using a Leica SPE-II equipped with a 63x oil immersion objective. Stacks of ~80 μm were acquired at 42.7 nm in *xy* and 420 nm in *z*. Staining of Cep192 lead to high background noise. The specific staining was therefore validated by co-localization of the alpha-tubulin staining or, in case of cold treated zygotes, it was replaced with NEDD1, which showed minimum background staining. To exclude that dual spindles resulted from polyspermy, we stained for acetylated tubulin of the residual sperm flagellum and only documented and analyzed embryos with a single flagellum (Fig. S1A) or scored for diploidy comparing the volumes of segmented DNA.

### Expression constructs and mRNA synthesis

Constructs used in this study to synthesize mRNA were previously described: pGEMHE-H2B-mCherry (Kitajima et al., 2011), pGEMHE-EGFP-MAP4 (Schuh and Ellenberg, 2007). To generate pGEMHE-EB3-EGFP2, full length *homo sapiens* EB3 cDNA (NM_001303050.1, a generous gift from Niels Galjart) was tagged at the C-terminus with a tandem mEGFP and cloned into the vector pGEMHE with a T7 promotor sequence for mRNA production. From linearized template DNA (1 μg), capped and poly-adenylated mRNA was synthesized *in vitro* using the mMESSAGE mMACHINE™ T7 ULTRA Transcription kit (AM1345, ThermoFisher Scientific). The mRNA was purified (74104, RNeasy Mini Kit, QUIAGEN) and dissolved in 14 μl RNAse free water.

### Micromanipulation

The cow zygotes were injected with mRNA in solution as described for mouse oocytes (Schuh and Ellenberg, 2007; Jaffe and Terasaki, 2004) with some modifications: In brief, a wider ‘injection slit’ was created between two glass cover slips by using a spacer of two layers of double sided adhesive tape (05338, tesa) tightly pressed together (~170-190 μm) to accommodate the cow zygotes of ~120 μm in diameter. The coverslip with the injection slit was attached to the plastic support slide with silicone grease and the whole chamber filled with 37-38 °C warm MOPS buffer before pipetting the embryos into the slit. The injection volume (4-5 pl) was adjusted to ~0.5% of the bovine zygotic volume. The mRNA concentrations ranged between 0.1-0.2 μg/μl for H2B-mCherry, 0.5 or 0.9 μg/μl for EB3-mEGFP2 and 0.3-0.4 μg/μl for MAP4-EGFP.

### Live imaging

For time-lapse imaging of cow zygotes, the in-house built inverted light-sheet microscope was used (Strnad et al., 2016) with the modifications described previously (Reichmann et al., 2018a), but using a 25 μm thick FEP foil (Lohmann, RD-FEP100A-610) as transparent base for sample mounting that allowed for easier handling due to reduced rigidity. In this setup the pixel size in *xy* 130 nm. For imaging of chromatin and either microtubule tips or lattice, fluorescence from H2B-mCherry and either EB3-mEGFP2 or EGFP-MAP4 was acquired simultaneously every 2.5 minutes using a 488 nm laser (~25-30 μW) and a 561 nm laser (~5-10 μW) with an exposure time of 100 ms. Stacks of 100-104 μm were acquired by 101 planes, resulting in a *z*-step size of 1-1.04 μm.

### Image processing

Time-lapse images were processed to extract single color data from the raw camera data as described previously (Strnad et al., 2016). We also used Fiji (Schindelin et al., 2012) with a new in-house built plugin for processing of large image data (Tischer et al., 2020).

### Quantification of alpha-tubulin IF intensity

An in-house developed MATLAB (The MathWorks, Inc.) script was utilized to quantify IF intensity from alpha-tubulin staining and perform a robust comparison of the intensity distributions of one spindle half with respect to the other. The script first segmented the signal from the metaphase chromosomes from the separate Hoechst channel and predicted the orthogonal spindle axis from the shape of the metaphase chromosomes. It then generated a set of parallel and equidistant cross sections of the tubulin channel orthogonal to the predicted axis.

To segment chromosomes, the Hoechst channel was first interpolated along the z direction to generate an isotropic 3D stack from anisotropic raw data. A 3D Gaussian filter was applied on the interpolated stack to reduce the noise where sigma and kernel size of the filter were set to 2 and 3, respectively. The Hoechst channel was binarized by combining parameters from adaptive thresholding (Otsu, 1979) applied on individual xy-planes of a z-stack, as well as on all xy-planes of the stack together (Hériché et al., 2014). The chromosome mass was identified by connected component analysis of the detected binary objects followed by smoothing operations. The spindle region was also detected using a similar approach, while centrosome coordinates were picked manually. 3D coordinates of all the voxels belonging to the detected chromosome mass were used to construct a Hessian matrix. The Eigenvector with the lowest Eigenvalue of this matrix approximates an orthogonal vector to the metaphase plate and thus was taken as the predicted spindle axis.

The predicted spindle axis was used as a reference to slice the microtubule channel at 500 nm spacing generating a set of parallel cross sections orthogonal to this axis. The slicing procedure was described in detail in Walther et al (Walther et al., 2018). A total of 24 μm in length along the predicted axis-12 μm in each direction from the centroid of chromosome mass – was taken for slicing. The size of a slice was 621×621 pixels, where the cross-section with the predicted spindle axis defined the center of the slice. To quantify microtubule intensity, the average background intensity was estimated first and was subtracted from the tubulin channel before the slicing. The average background intensity was calculated from a rim of 2 pixels in width with a distance of 8 pixels from the boundary of the segmented spindle region. The total background subtracted intensity of tubulin inside each slice was plotted with respect to its distance from the centroid of the chromosome mass (see Fig. 2B, Fig. S2A-B). This intensity profile was further analyzed to extract different parameters to describe the shape and intensity of two spindle halves:

The valley between two intensity peaks from the profile was detected first to define parts of the profile belonging to individual spindle halves. Total intensity of a spindle half was calculated by summing up the area under the intensity profile curve (AUC) belonging to that spindle half. To estimate the width of intensity distribution belonging to a spindle half, the intensity at the valley was subtracted from the profile (opaque area under distribution valley, Fig. 2B). Full width half maxima (FWHM) of the valley subtracted intensity distribution from each spindle half were calculated. Both parameters (total intensities and FWHMs of spindle halves) were normalized to either the acentrosomal half (for monocentrosomal spindles) or to the half with the smaller measure (for bicentrosomal spindles) (Fig. 2C-D).

The length of a spindle half was also calculated from the respective part of the original intensity profile (without subtracting the intensity at the valley). The distance between the valley and the other edge of the intensity distribution, representing the slice at the periphery of a spindle half, was used to determine the length. The ‘polar’ periphery of each spindle half was detected by taking the peripheral position closest to the peak of the profile where intensity value was less than 10% of the peak. If no such slice was found (all values are higher than 10% of the maximum, n = 1), the furthest slice from the peak was considered as periphery of a half. The lengths of the spindle halves were also normalized to the either the acentrosomal half (for monocentrosomal spindles) or to the shorter half (for bicentrosomal spindles) (Fig. 2E).

### Calculating distances from centrosomes to the spindle body and to chromosomes

To calculate the distance between centrosome and spindle body (d_1_, Fig. S1E), the axis between the centrosome and chromosome centroid was used as a reference for slicing the microtubule channel (slicing described in previous section). In this case slicing along this axis was performed at 200 nm spacing starting from the centrosome towards the chromosome centroid. The total intensity of each slice was calculated in order to create an intensity profile along the axis. The maximum and minimum intensity values were determined first and the minimum intensity was subtracted from the profile. The minimum subtracted profile was probed starting from the centrosomal end. The last location in the profile, where the total intensity was less than 10% of the maximum total intensity, was determined as the periphery of the spindle body. The distances between the manually annotated centrosome coordinates and the determined spindle body (d_1_) or the chromosome centroid (d_3_) and between the chromosome centroid and the spindle body (d_2_; Fig. S1E) were then calculated (Fig. S1F and S1H).

This centrosome-spindle distance (d_1_; Fig. S1E) was normalized to the length of the respective spindle half (d_2_) to illustrate distance relations (d_1_/d_2_; Fig. S1G).

### Quantification of dynamic microtubule distribution of EGFP-MAP4 and EB3-mEGFP2 signal

Microtubule signal intensity (EGFP-MAP4 or EB3-mEGFP2) was quantified along the centrosomal axis defined by the two centrosomes. The center of intensities and thus central coordinates of the centrosomes were determined from manually segmented microtubule signal at centrosomes using arivis Vision4D. Original anisotropic stacks were first interpolated along the *z* direction to create isotropic 3D stacks and a microtubule intensity profile from one centrosome to the other was generated. For each time point a total of 15 equidistant parallel slices starting from one centrosome to the other were generated. The slices were taken orthogonal to the centrosomal axis where the center of each individual slice was intersected by the axis. The size of a slice was set to either 351×351 pixels (black cuboid with solid line, Fig. 4B) to determine the total microtubule mass along the centrosomal axis and within the entire spindle at a given time (Fig. 4C). Or the slice size was set to 15×15 pixels (black cuboid with dashed line, Fig. 4B) to estimate the microtubule concentration along the centrosomal axis (Fig. 4D). The distance between two centrosomes was variable in different time points within a zygote as well as between zygotes. To address this, a fixed number of slices between two centrosomes were generated. This normalized the distances between two consecutive slices in different stacks with respect to the distance between centrosomes. The total intensity of each slice was calculated and was normalized to the maximum total intensity considering all slices and all time points within a zygote. Normalization of intensity and inter-slice distance made the extracted intensity profile comparable within a zygote as well as between different zygotes. This allowed computation of an average intensity profile (Fig. 4C and D) over time and intensity change over time at different landmarks (such as centrosomes and the center of the spindle, see Fig. 4E and F) using the data from all the analyzed zygotes (n = 6).

### Data transformation into 2D sections parallel to the centrosomal axis

To display the kinetics of microtubule intensity over time (Fig. 4A), 3D data was transformed and re-sliced orthogonal to the centrosomal axis so that both centrosomes were visible in the same 2D slice. The raw data was interpolated as described in the previous section to generate an isotropic 3D stack. The interpolated data was then translated in *xy* so that the midpoint between two centrosomes moved to the center of the translated image. The angle between centrosomal axis (defined by the two centrosomes), and xy-plane was calculated and the translated stack was rotated to align the centrosomal axis to the xy-plane. The angle between centrosomal axis and x axis was calculated and the data was further rotated to align the centrosomal axis to the original x axis. Bicubic interpolation was used during the rotation. All the data was transformed in the same way so that the kinetics of microtubule intensity at centrosomes as well as the center of the centrosome axis could be observed in the same 2D slice over time.

## Acknowledgements

Confocal microscopy images were acquired at the Centre for Cellular Imaging at the Faculty of Veterinary Medicine, Utrecht. The authors would like to thank Richard Wubbolts and Esther van ‘t Veld from the Centre for Cellular Imaging for their help and technical assistance in image acquisition. We also thank Christian Tischer from the Center for Bioimage Analysis at EMBL for vital support with automated image processing and Petr Strnad for the development of the inverted light-sheet microscope and Yu Lin, Lars Hufnagel and Balint Balazs for maintenance and software modifications. The authors would like to thank Claudia Deelen, Radyon Huggins, Anouk Klein Kranenbarg and Romy Timmer for their assistance with IVF and IF staining. The authors would also like to thank Andrea Genthner and Klaus Schmitt for assistance on the transport and work with bovine COCs at EMBL Heidelberg. The authors also thank Nathalie Daigle and Tomoya Kitajima for cloning of the DNA constructs that were used as templates for mRNA synthesis in this study. We thank Judith Reichmann and Manuel Eguren for helpful scientific discussions, and we also thank the whole Ellenberg Lab at EMBL and the IVF Lab of the Faculty of Veterinary Medicine in Utrecht for collegial support. We thank Franziska Kundel for reading the manuscript and Stephanie Alexander for organizational support. The work was supported by funds from the European Research Council (ERC advanced Grant,,COREMA”, grant agreement 694236) to Jan Ellenberg and the European Molecular Biology Laboratory. Isabell Schneider was further supported by a Boehringer Ingelheim Fonds PhD fellowship and Marta de Ruijter-Villani by an EMBO short-term fellowship for this project. Competing interests: Jan Ellenberg is scientific co-founder and advisor of Luxendo GmbH (part of Bruker), which makes light-sheet-based microscopes commercially available. The authors have no additional competing financial interests.

## Author contributions

Jan Ellenberg, Marta de Ruijter-Villani and Isabell Schneider conceived the project. Isabell Schneider and Marta de Ruijter-Villani further designed and conducted the experiments and wrote the original draft of the manuscript. M. Julius Hossain, Isabell Schneider and Marta de Ruijter-Villani performed the formal analysis. Tom A. E. Stout contributed to conception of the work and edited the manuscript. Jan Ellenberg supervised the project and reviewed and edited the manuscript. All authors contributed to the interpretation of the data and read and approved the final manuscript.

## Abbreviations

AUC: area under the curve
IF: immunofluorescence
IVF: in vitro fertilization
FWHM: full width half maximum
MTOC: microtubule organizing center
NEBD: nuclear envelope break down
PN/s: pronucleus/pronuclei

## Supplementary figure legends

**Figure S1:**
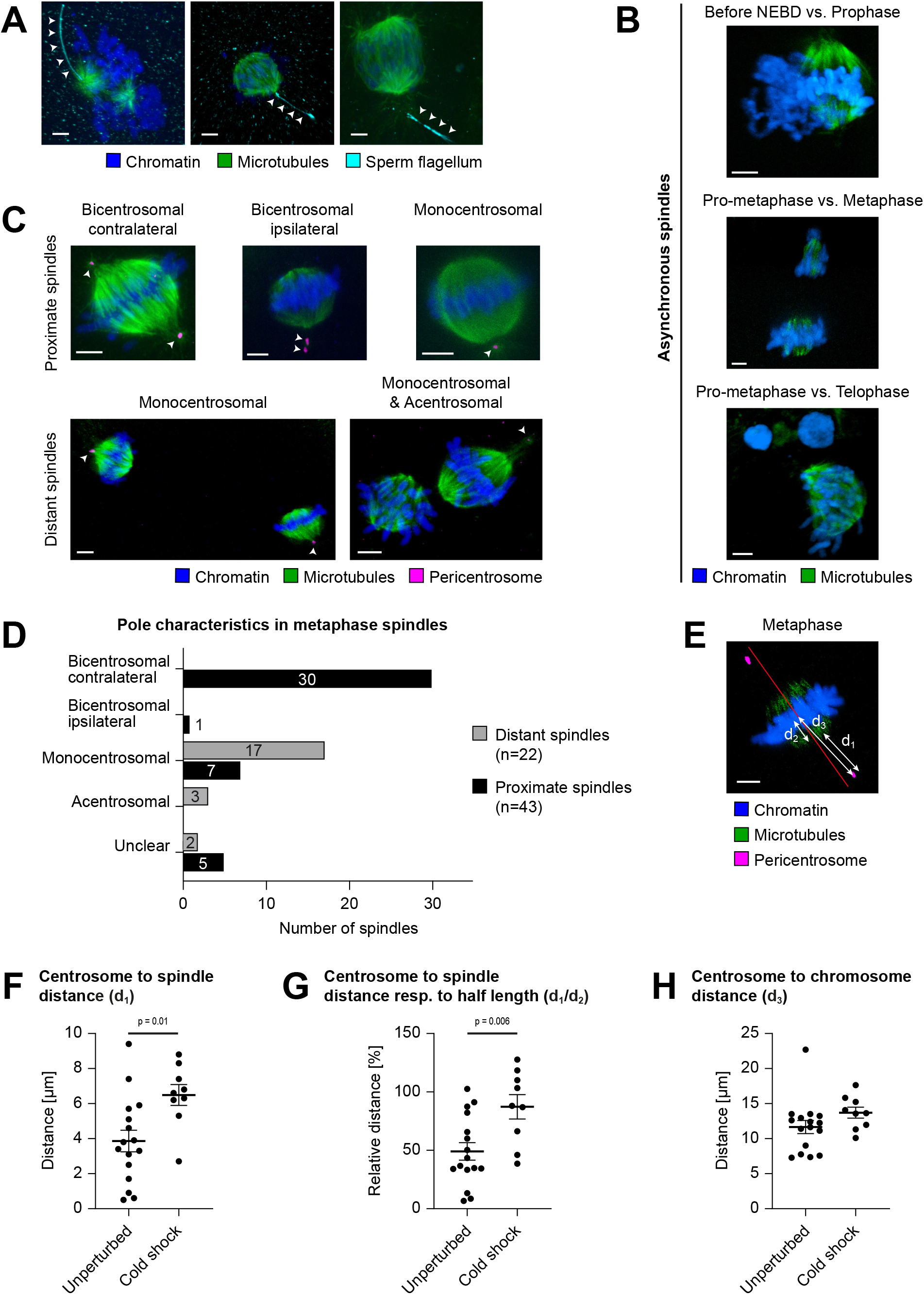
Dual spindle characteristics in bovine zygotes. **(A-C, E)** Immunofluorescence staining of bovine zygotes fixed at 27.5 h post *in vitro* fertilization. Maximum intensity projections orthogonal to the spindle axis of confocal sections are shown. Microtubules (alpha-tubulin, green); chromatin (Hoechst, blue), sperm flagellum (acetyl tubulin, cyan), pericentrosome (Nedd1, Cep192, magenta). Scale bars, 5 μm. **(A)** White arrow heads indicate spermatozoan flagellum adjacent to one spindle pole confirming monospermic fertilization. For further details, see Materials and Methods. **(B)** Distant dual spindles with distinct mitotic timing inside same cytoplasm. **(C-D)** Diverse centrosome positioning in proximate (fused) and distant dual spindles. **(C)** Arrowheads indicate the number and position of centrosomes. **(D)** Abundance of centrosome positions as illustrated in (C). **(E-H)** Comparison of centrosome positioning in bicentrosomal (contralateral) metaphase spindles after IF of unperturbed (n = 16) or cold-treated (3 min cold shock on ice, n = 9) zygotes. d_1_: centrosome to spindle distance; d_2_: spindle half length; d_3_: centrosome to chromosome distance. **(E)** Illustration of the measurements in exemplary metaphase spindle after cold treatment (see also Fig. 1D), red line illustrates projected spindle axis orthogonal to chromosomes. **(F)** Distance in μm between centrosomes and spindle body (d_1_, see arrow in E). For assessment of spindle body, see Material and Methods. Bars indicate standard error of the mean distance of unperturbed (3.9 μm) vs. cold shock treated (6.5 μm) zygotes (p = 0.01, significant). **(G)** Relative distance between centrosome and spindle microtubules respective to spindle half length (d_1_/d_2_, see arrows in E). Bars indicate standard error of the mean distance in unperturbed (49.1%) vs. cold shock treated (87.3%) zygotes (p = 0.006, significant). **(H)** Distance in μm between centrosomes and the chromosome centroid (d_3_, see arrow in E). Bars indicate standard error of the mean distance in unperturbed (11.7 μm) vs. cold shock treated (13.7 μm) zygotes (p = 0.15). (F-H) Average measurements for both centrosomes from same zygote are depicted. Statistical test: Unpaired t-test.

**Figure S2:**
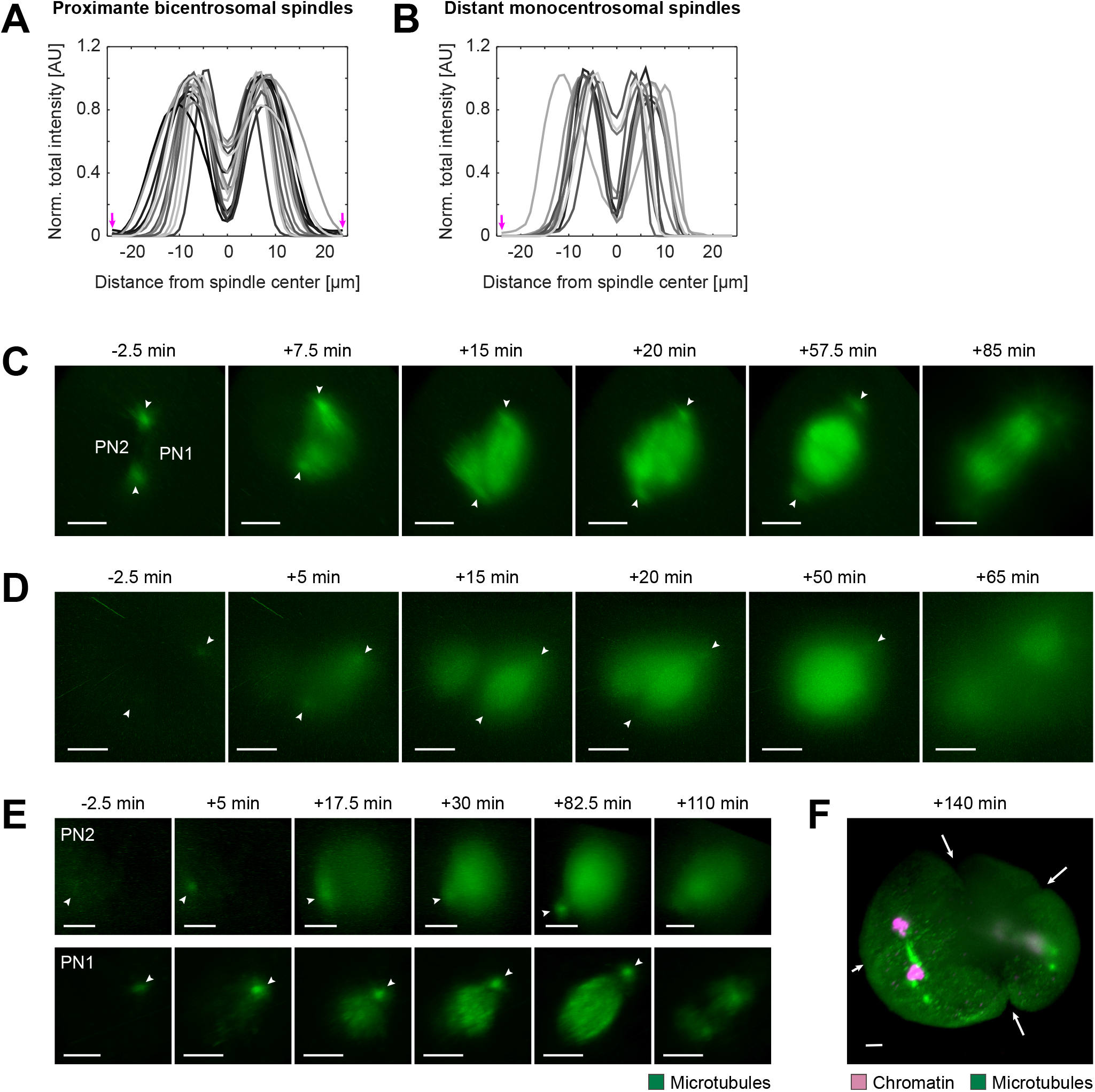
Comparing microtubule distribution in proximate bicentrosomal and distant monocentrosomal spindles in fixed and live bovine zygotes. **(A-B)** Intensity distribution of alpha-tubulin immunofluorescence in 2D sections along the calculated spindle axis orthogonal to the metaphase chromosomes in both, proximate bicentrosomal contralateral (A) and distant monocentrosomal (B) spindles (see also Fig 2A-B). Magenta arrows indicate positions of centrosomes. **(C-E)** Respective 3D rendered images of fluorescence from microtubule markers (EGFP-MAP4 and EB3-mEGFP2) in the pronuclear volumes of zygotes shown in Fig 3A-C to highlight dual spindles, and centrosome positions (white arrow heads). Timings respective to synchronous pronuclear envelope breakdown (NEBD) or to NEBD of the first pronucleus (PN1) in case of asynchrony. PN2, lagging pronucleus. Projected scale bars, 10 μm. **(F)** 3D rendered image of fluorescence from microtubule marker (EB3-mEGFP2, green) and chromatin marker (H2B-mCherry, magenta) of zygotic volume (same zygote as Fig 3C, S2D and E) after background correction (median based denoising). White arrows indicate multiple ingression sites at 140 min post NEBD as consequence of distant dual spindles.

**Figure S3:**
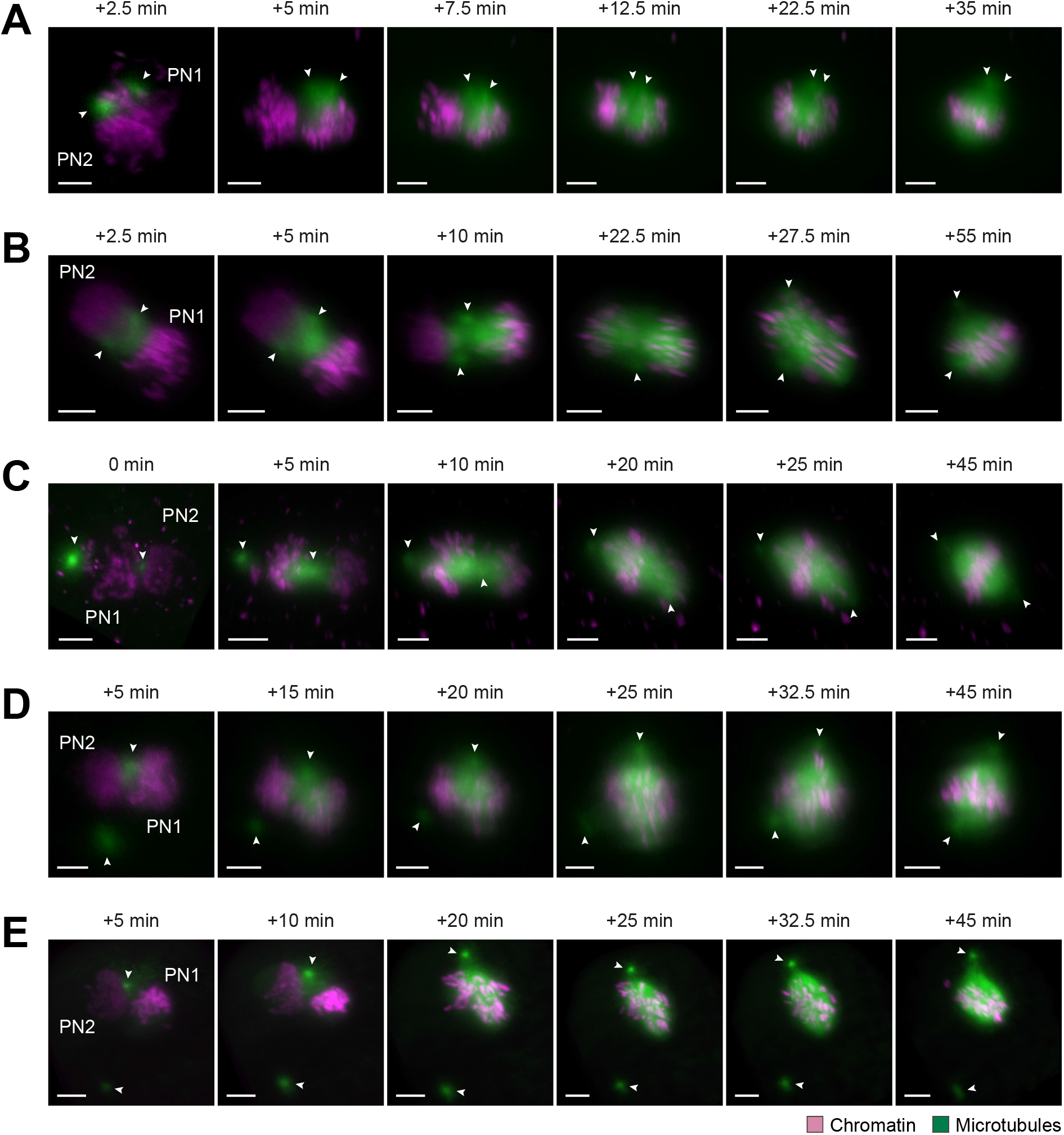
Miscellaneous spindle assembly modes around proximate parental genomes in live bovine zygotes. **(A-E)** Bovine zygotes expressing microtubule markers (EGFP-MAP4, A-D or EB3-mEGFP2, E; green) and chromatin marker (H2B-mCherry; magenta) were imaged by light sheet microscopy every 2.5 min throughout mitosis and for up to 6 h in total. 3D rendered images of pronuclear volumes after background correction (median based denoising) show examples of spindle formation and dynamics from Pro-to Metaphase. Indicated timings respective to synchronous pronuclear envelope breakdown (NEBD) or to NEBD of first pronucleus (PN1) in case of asynchrony. PN2; lagging pronucleus. Arrow heads indicate positions of centrosomes. Projected scale bars, 10 μm. **(A-C)** Spindle assembly modes around adjacent pronuclei (PNs), where centrosomes localized at PN surfaces, but not at the PN interphase junctions (n = 4). Centrosomes either localized in proximity to each other but not perfectly at PN interphase (A and B) or at opposite sides of same PN, with only one centrosome at PN interphase (C). **(D-E)** Spindle assembly around adjacent PNs, where only one centrosome localized to PN surface/interphase and the second was randomly positioned in cytoplasm without clear nuclear attachment (n = 3).

## Supplementary movie legends

Movie S1: Time-lapse imaging of mitotic live bovine zygote expressing EGFP-MAP4 (green) and H2B-mCherry (magenta) after mRNA injection at pronuclear stage. Time resolution; 2.5 min. Scale bar, 10 μm. Movie shows 30 frames/s. Recording starts at 2.5 min prior to NEBD of the leading pronucleus (PN1). Movie shows spindle assembly in zygote depicted in Fig 3A and S2C. It is also an example used for analysis, see Fig 4C-F.

Movie S2: Time-lapse imaging as in Movie S1, but of zygote expressing EB3-mEGFP2 (green) and H2B-mCherry (magenta). Recording starts at 2.5 min prior to synchronous NEBD. Movie shows spindle assembly in zygote depicted in Fig 3B and S2D.

Movie S3-5: Time-lapse imaging of mitotic live bovine zygote as in Movie S1. Recordings start at 2.5 min prior to NEBD of the leading pronucleus (PN1). Movies show spindle assembly around the distant pronuclear volumes (PN2 and PN1, Movie S3 and 4, respectively) or in the context of the entire imaged volume (Movie S5) from same zygote, also depicted in Fig 3C and S2E-F.

